# Guidelines for standardising the application of discriminant analysis of principal components to genotype data

**DOI:** 10.1101/2022.04.13.488270

**Authors:** Joshua A. Thia

## Abstract

Discriminant analysis of principal components (DAPC) has become a popular method for visualising population structure due to its simplicity, computational speed, and freedom from demographic assumptions. Despite the popularity of DAPC, there has been little discussion on best practise. In this work, I provide guidelines for standardising the use of DAPC in studies of population genetic structure. An often-overlooked fact is that DAPC generates a model describing the genetic differences among a set of populations defined by a researcher. I demonstrate that appropriate parameterisation of this model is critical for obtaining biologically meaningful results. I show that the number of leading PC axes used as predictors of among population differences, *p*_axes_, should not exceed the *k* – 1 biologically informative PC axes that are expected for *k* effective populations in a genotype dataset. This *k* – 1 criterion for *p*_axes_ selection is more appropriate compared to the widely used *proportional variance criterion,* which often results in a choice of *p*_axes_ ≫ *k* – 1. DAPC parameterised with no more than the leading *k* – 1 PC axes is: (1) more parsimonious; (2) captures maximal among-population variation on biologically relevant predictors; (3) less sensitive to unintended interpretations of population structure; and (4) more generally applicable to independent sample sets. Assessing model fit should be routine practise and can aid interpretation of population structure when implementing DAPC. Additionally, it is imperative that researchers clearly articulate their study goals, that is, testing *a priori* expectations versus studying *de novo* inferred populations. Distinguishing between these goals is important because it dictates whether a researcher’s results can be treated as a test of the hypothesis that significant genetic differences exist among populations. Defining populations *a priori* (before observing the genotype data) constitutes a true hypothesis test, but populations defined *de novo* (after observing the genotype data) cannot be used to test this hypothesis due to issues with circularity. The discussion and practical recommendations provided in this work provide the molecular ecology community a roadmap for applying DAPC to their genotype datasets.

## Introduction

The biological world is beautifully complex, characterised by variation in multiple dimensions. Multivariate statistics play a pivotal role in helping us make sense of this multi-dimensionality and developing a deeper appreciation of biology. Describing population genetic patterns, for example, becomes increasingly difficult with many sampled individuals, genetic markers, and populations. However, ordination methods can summarise variation across multiple loci to create new synthetic axes and reduce dimensionality. Such new axes of variation facilitate the visualisation of genetic relationships among individuals and populations, aiding the interpretation of population structure.

Principal component analysis (PCA) is an incredibly useful multivariate method for summarising genetic differences among individuals. PCA reveals structure in a dataset by capturing the major axes of covariance among a set of *p* variables (Rencher, 2002c). Putative groupings can be inferred from the clustering of samples in discrete regions of PC space. Because PCA does not make any explicit assumptions about the organisation of variation in the *p* variables, PCA is a *hypothesis free* method, providing information about the innate structure in a sample (Rencher, 2002c). All PC axes are orthogonal and are constrained to capture decreasing amounts of variance from the first to last axis (Rencher, 2002c). Hence, in a structured set of variables, dimension reduction can be achieved by selecting some number of the leading PC axes that describe the most variation (Rencher, 2002c). The dimensionality reduction achieved by a PCA is hugely beneficial in population genomic studies, summarising the covariances among thousands, maybe millions, of loci (Patterson, Price, & Reich, 2006). Determining which PC axes capture biologically informative variation is a non-trivial task (Cattell, 1966; Jackson, 1993). However, for a genotype dataset comprising *k* effective populations, prior theoretical work indicates that only the first *k* – 1 PC axes capture population structure (Patterson et al., 2006).

Discriminant analysis (DA) is another multivariate method that has become popular in population genetic studies through the discriminant analysis of principal components (DAPC) approach (Jombart, Devillard, & Balloux, 2010) (Figure 1). DA takes a set of measured *p* predictor variables and identifies linear combinations of those variables that maximise discrimination of individuals from defined groups. Because DA models the differences among groups in their multivariate means, DA is a *hypothesis driven* method (Rencher, 2002a), and is related to a multivariate analysis of variance (Rencher, 2002b). Performing DA directly on a genotype dataset of many loci is undesirable because the number of *p* variables (genetic loci) often greatly exceed the number of *n* sampled individuals, and there may be extensive correlations among loci (Jombart et al., 2010). The DAPC approach circumvents this issue by first performing a PCA to reduce dimensionality and remove correlation, with the leading p_axes_ number of PC axes used as predictor variables to discriminate among *k*_DA_ groups in a DA. The value of *k*_DA_ is chosen in one of two ways: (1) *k*_DA_ = *k*_prior_, where *k*_prior_ is an *a prior* expectation about the number of populations, for example, the number of sampling locations; or (2) *k*_DA_= *k*_infer_, where *k*_infer_ is the *de novo* designation of populations, an inference of the *k* number of effective populations obtained from genotype data (as per descriptions in Miller, Cullingham, & Peery, 2020). (Note, relevant mathematical notations are detailed in Table 1.)

**Figure 1.**
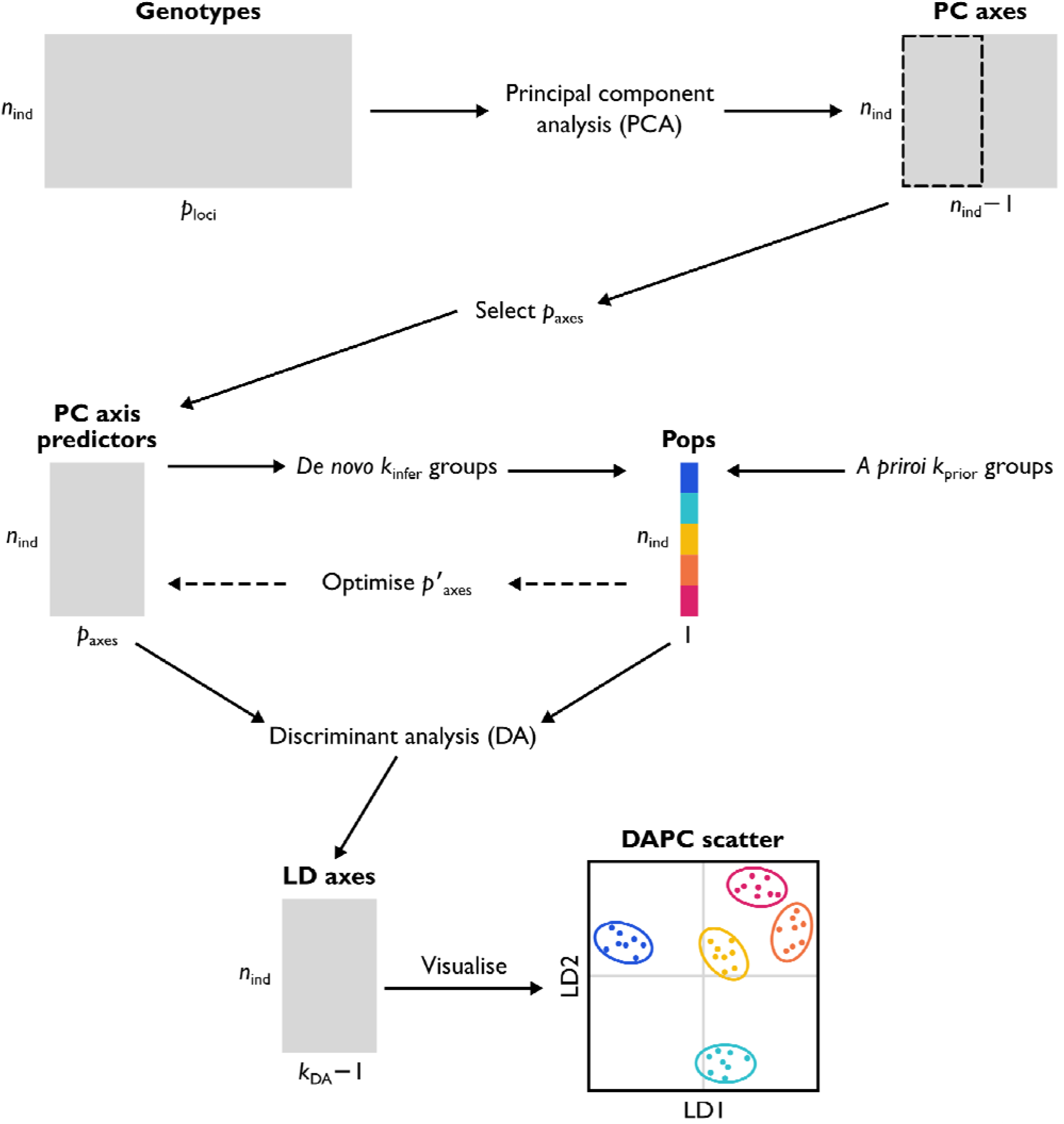
Schematic representation of a DAPC (discriminant analysis of principal components). A genotype matrix comprising *n*_ind_ samples genotyped at *p*_loci_ genetic loci is subjected to a PCA (principal component analysis). Assuming that *n*_ind_ < *p*_loci_, there are *n*_ind_ – 1 PC axes with non-zero eigenvalues that summarise the allelic covariances in the sample. A subset of these PC axes, *p*_axes_, are selected to represent population structure. Grouping of individuals into populations may be based on some *a pirori* expectation, *k*_prior_, or may be inferred *de novo* from the PCA, *k*_infer_. Optionally, the *adegenet* functions xvalDapc or optim.a.score can be used to produce an optimised number of predicted PC axes, *p*’_axes_. DA (discriminant analysis) is then used to identify linear combinations of the PC axes as predictors of differences among *k*_DA_ groups, where *k*_DA_ = *k*_piror_ or *k*_DA_ = *k*_infer_. Assuming *p*_axes_ ≥ *k*_DA_ – 1, a set of *k*_DA_ – 1 LD (linear discriminant) axes is returned that summarise the among population variation. Visualisation of individuals projected into LD space is used to infer genetic relationships among individuals and populations.

**Table 1.** Summary of mathematical notation.

| Notation | Definition |
| --- | --- |
| $k$ | The true number of effective populations in a genotype dataset. |
| $k_{\text{prior}}$ | The *a priori* expectation of the number of populations, based on criteria external to the genotype data, for example, sampling locations. Sampled individuals are designated to one of these *a priori* defined populations. |
| $k_{\text{infer}}$ | The inferred number of effective populations, an estimate of $k$, from genotype data. Can be used to guide *de novo* population designation of sampled individuals. |
| $k_{\text{cluster}}$ | The number of clusters to divide a sample among in *K*-means clustering. Many values could be tested. The most likely value of $k_{\text{cluster}}$ can be used as inference of the number of effective populations, $k_{\text{infer}}$. |
| $k_{\text{DA}}$ | The number of groups to discriminate among in a DAPC, a researcher's choice of $k_{\text{prior}}$ or $k_{\text{infer}}$ to analyse. |
| $p_{\text{axes}}$ | The number of leading PC axes to use as predictors of differences among $k_{\text{DA}}$ groups in a DAPC. |
| $p'_{\text{axes}}$ | The optimised number of leading PC axes to use as predictors of differences among $k_{\text{DA}}$ groups in a DAPC. |
| $m$ | The individual migration rate from a source population into a focal population. |
| $M$ | The total migration rate into a focal population, the summed $m$ across all source populations. |

Despite the frequent use of DAPC by molecular ecologists, best practise for this approach has barely been discussed. Such matters were recently addressed in a combined meta-analysis and simulation study by Miller et al. (2020). In their work, Miller et al. (2020) found that many studies fail to report the relevant parameters required to replicate their DAPC results. They also provided some practical suggestions on how DAPC of genotype data can be made more transparent by reporting how groups are defined, clearly specifying the goals of a DAPC, and stating the number of PC axes used in the analysis. Miller et al. (2020) highlighted that the molecular ecology community has not yet converged on set of standard operating procedures, which leads to variable parameterisation among studies. Missing from the literature is clear communication of how DAPC functions “under the hood” and the necessary considerations required by researchers when using this approach to study population structure.

In this piece, I address the need for best practise guidelines to standardise the application of DAPC to genotype data. Researchers should be aware that when performing a DAPC they are fitting a model that describes the genetic differences among defined groups. This model is produced by the DA portion of a DAPC, which uses genotypic PC axes as predictors of among-population genetic differences. As I demonstrate here, careful parameterisation of this model is critical in obtaining biologically relevant results. I show that the appropriate number of PC axes for DA is a deterministic property of a genotype dataset: the leading *k* – 1 PC axes for *k* effective populations. I propose a *k* – 1 criterion for choosing a value of *p*_axes_ and argue that this criterion is more appropriate for DAPC parameterisation than the commonly used *proportional variance criterion*, where *p*_axes_ is instead chosen to capture some proportion of the total genotypic variance. The fit of the model produced by DA should be assessed through methods such as leave-one-out cross-validation or training-testing partitioning. Furthermore, I explain the importance of clearly defining the goals of a DAPC (testing *a priori* expectations, or studying *de novo* inferred populations) because this has implication as to how researchers should interpret their results.

## Simulated population genetic datasets

To investigate how parameterisation choices affect the results and downstream interpretation of DAPC, I simulated metapopulations connected by different rates of gene flow. These metapopulations comprised five constituent populations and followed finite island model dynamics (Takahata & Nei, 1984; Wright, 1931). All populations were the same size and underwent divergence with or without gene flow. Simulations were performed using *fastsimcoal2* v2.7 (Excoffier, Dupanloup, Huerta-Sánchez, Sousa, & Foll, 2013; Excoffier et al., 2021). Each population was comprised of 1,000 haploid genomes (500 diploids). Each genome was 20 Mb in length with a recombination rate between adjacent bases of 1e–8 and a mutation rate of 2e-9. Populations underwent a series of hierarchical divergence events, with each divergence event 10,000 generations from the next (Figure S1). I modelled three migration scenarios with different rates of gene flow, with *M* as the overall rate of gene flow into a focal population, and *m* as the individual contribution of each source population into a focal population: (a) *M* = 0.000, *m* = 0.000, divergence with no gene flow, *F*_ST_ = 0.99; (b) *M* = 0.004, *m* = 0.001, divergence with gene flow, *F*_ST_ = 0.09; and (c) *M* = 0.400, *m* = 0.100, an effectively panmictic (homogeneous) metapopulation, *F*_ST_ = 0.0009 (see Figures S2 and S3 for simulation summaries). These scenarios reflect different contexts where the number of perceived sampled populations may or may not be equal to the number of *k* effective populations. In the *M* = 0.000 scenario, there is strong population structure, and a clear *k* = 5. In the *M* = 0.004 scenario, population structure is weaker, but there is still a discernible *k* = 5. In the *M* = 0.400 scenario, there is no population structure, and *k* = 1.

For each migration scenario, I created two distinct simulated datasets. The first involved a single simulation at each migration rate where all 500 diploid contemporaneous individuals per population were sampled at the end of the simulation: herein, the *singular simulation dataset.* The purpose of this first dataset was to illustrate variation and relationships among independent sample sets from the same simulated metapopulation. Diploid individuals were split into 20 sample sets, each containing 25 individuals per population. In the second dataset, I performed 30 independent simulations at each migration rate and sampled 25 diploid contemporaneous individuals per population at the end of the simulation: herein, the *replicate simulation dataset.* The purpose of this second dataset was to illustrate generalities across independent metapopulations that had undergone the same migration scenario.

Simulation results were imported into R v4.1.3 (R Core Team, 2022). For each simulation in the *replicate simulation dataset,* I filtered SNP loci to a sample size of 1,000 random SNPs distributed across the genome. First, the genome was divided into 5 Kb windows, from which one random SNP was sampled. Second, these remaining SNPs were randomly sampled to retain 1,000 loci. For the *singular simulation dataset,* a preliminary filtering was performed to remove all loci that were not polymorphic across all discrete sample sets. Following this, random sampling per genomic window and reduction to 1,000 random loci was performed. Details of all downstream analyses can be found in the Supplementary Information.

## Population structure captured by principal component analysis

Determining which PC axes, if any, capture significant covariance among measured variables presents a major challenge for the use of these dimensions in predictive analyses (Peres-Neto, Jackson, & Somers, 2005). Patterson et al. (2006) demonstrated a clear expectation that for *k* effective populations there are *k* – 1 PC axes that capture variation among populations, with all remaining axes characterising variation within populations. The inference of *k*, *k*_infer_, is obtained from the genotype data. In this section, I will highlight simple ways that *k*_infer_ can be obtained visually and explain why we should only focus on the leading *k* – 1 PC axes of population structure. Although various methods exist to statistically test the number of biologically informative PC axes (Jackson, 1993; Peres-Neto et al., 2005) and to obtain k_infer_ from genotype data (Patterson et al., 2006), I will not address the performance of these methods here. I will, however, discuss the use of *K*-means clustering to obtain *k*_infer_, as it is commonly used in conjunction with DAPC.

One simple way to visually obtain *k*_infer_ is by examining PC screeplots. Screeplots illustrate the decline in explained variance (or eigenvalues) associated with each sequential PC axis. When population structure exists, there is typically an inflection point at the *k*^th^ PC axis that creates an “elbow shaped” pattern in the explained variances (Abegaz et al., 2019; Cattell, 1966), as illustrated in Figure 2a. We can see that in the *M* = 0.000 and *M* = 0.004 scenario, the explained variance rapidly decreases from PC axes 1 to 4 (*k* – 1), and then decreases more incrementally beyond PC axis 5 (k). The scree is also steeper between PC axes 1 to 4 in the *M* = 0.000 scenario because more variation is among populations (*F*_ST_ = 0.99), relative to that in the *M* = 0.004 scenario (*F*_ST_ = 0.09). Contrastingly, in the *M* = 0.400 scenario, although there were five sampled populations, there is no variation among them (*k* = 1, *F*_ST_ = 0.0009), and the scree slope exhibits a smooth decline.

**Figure 2.**
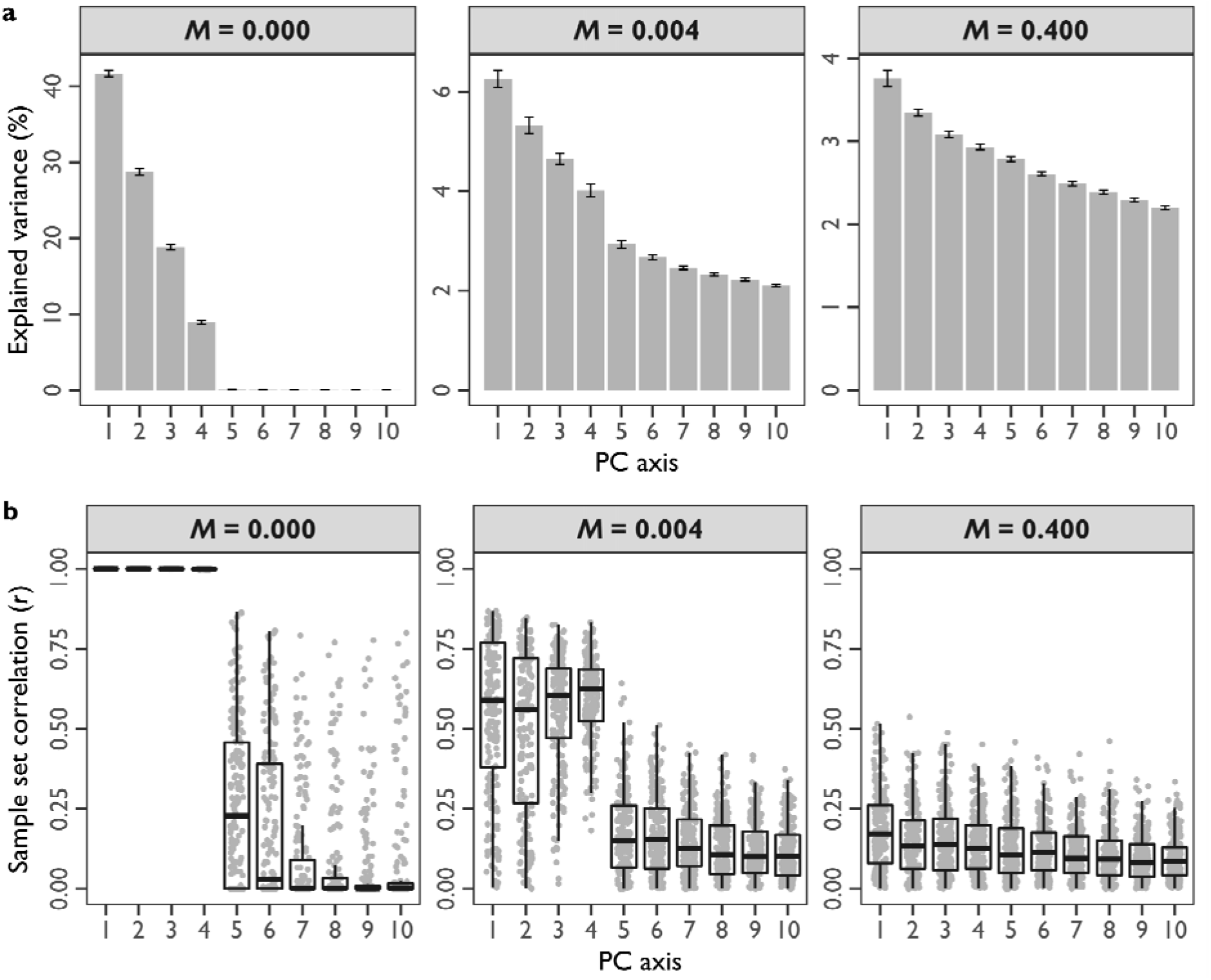
Principal component analysis under different migration scenarios, (a) Average explained variance for PC axes across simulated sample sets, using the *replicate simulation dataset*. The *x*-axis represents the first 10 PC axes. The *y*-axis is the amount of explained genotypic variance (percent of total) ascribed to each PC axis. Grey bars indicate the average across replicate simulations with the 95% confidence intervals, (b) Correlations among PC axis eigenvectors in the *singular simulation dataset.* The *x*-axis represents the first 10 PC axes. The *y*-axis represents the correlation of eigenvector loadings (contributions of each locus to variation on that PC axis). Each grey point represents a pair of independent sample sets from the same simulated metapopulation. Boxplots summarise the distribution of correlations across sample set pairs.

A second way to visually obtain *k*_infer_, and to validate inferences from screeplots, is by examining the scatter of individuals projected into PC space. Each dimension from PC axis 1 to *k* – 1 should capture different aspects of the among-population variation for *k* effective populations. PC axes ≥ *k,* however, are associated with stochastic sampling noise and within-population variation. Therefore, clustering of samples should only be observed on PC axes ≤ *k –* 1. These expectations are illustrated in Figure 3. For *M* = 0.000 on PC axes 1 to 4 (Figures 3a and 3b), individuals from each population pile up into very discrete regions of PC space because almost all the variation is among populations. For *M* = 0.004 on PC axes 1 to 4 (Figures 3d and 3e), population clusters are more dispersed, and their multivariate means are closer in PC space, because genetic differentiation is weaker. For both *M* = 0.000 and 0.004, beyond PC axis 5 (Figures 3c and 3f, respectively), all populations collapse into a single homogeneous data cloud. In contrast, for the *M* = 0.400 scenario, there is no clustering of individuals in PC space on the first four PC axes, and all the variation is within populations because *k* = 1.

**Figure 3.**
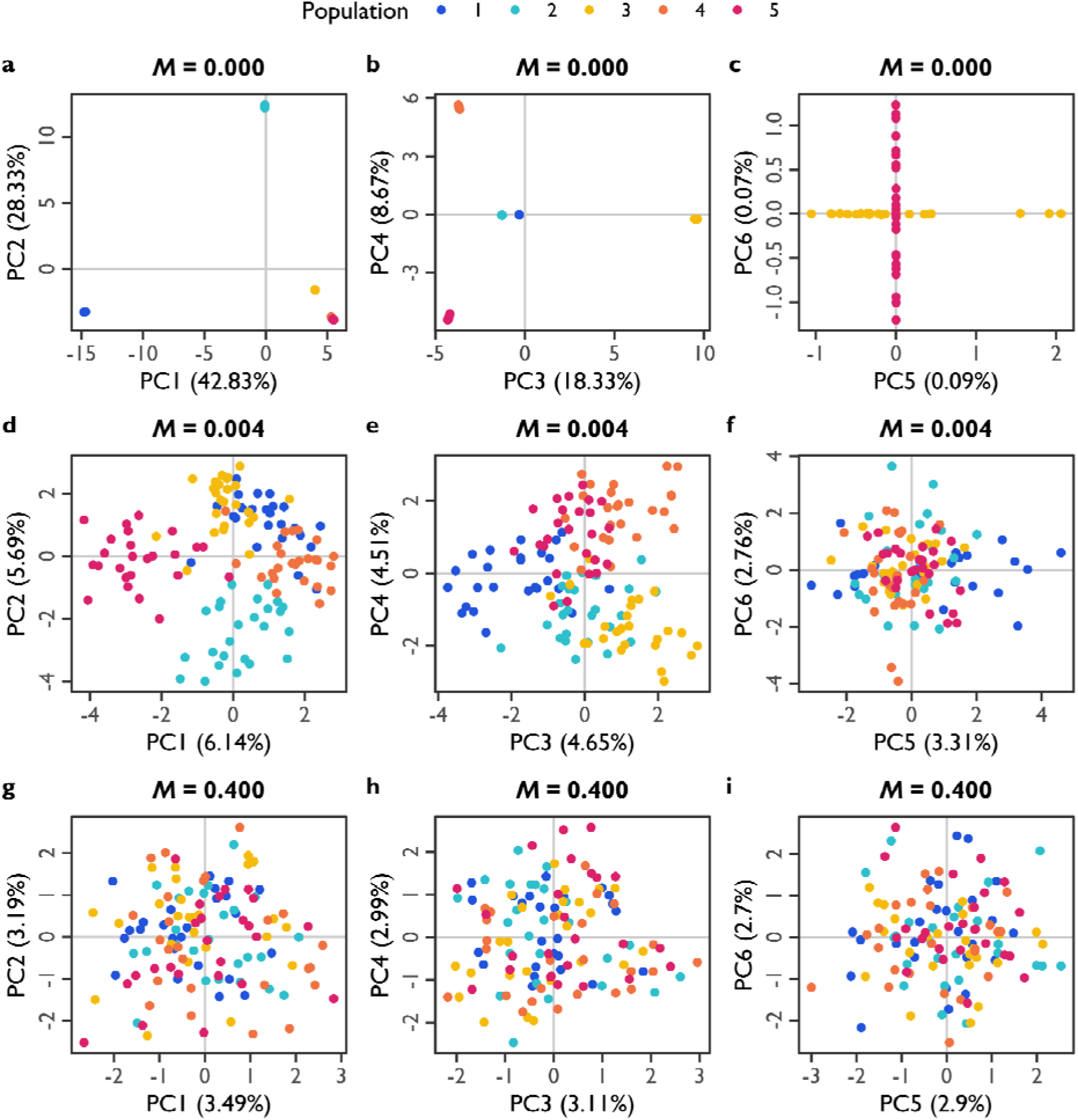
Exemplar scatterplots of individuals projected into PC space. The x-axis and y-axis, respectively, represent different pairs of PC axes. Numbers in parentheses indicate the amount of explained genotypic variance captured by each PC axis (percent of total). Each point represents an individual, coloured by population (see legend). Migration rates: (a, b, c) *M* = 0.000; (d, e, f) *M* = 0.004; and (g, h, i) *M* = 0.400. PC axis pairs: (a, d, g) 1 versus 2; (b, e, h) 3 versus 4; and (c, f, i) 5 versus 6.

*K*-means clustering offers another way to obtain *k*_infer_. The goal of *K*-means clustering is to allocate observations into a specified number of *k*_cluster_ clusters by minimising the within-group sum of squares (Hartigan and Wong, 1979). This method is commonly implemented within the DAPC pipeline, although it is not an essential element of DAPC. Note, that like DA, *K*-means clustering is also a *hypothesis driven* method. When performing *K*-means clustering, a researcher is testing the hypothesis that exactly *k*_cluster_ groups occur in their data. Many values of *k*_cluster_ can be fitted and compared using information criteria as the test statistic, for example, a BIC score. The most likely value can then be chosen to represent *k*_infer_, and to inform groups *de novo* for DA with *k*_DA_ = *k*_infer_. Predictors of the *k*_cluster_ groups are typically the same number of leading PC axes, *p*_axes_, that will be used in DAPC. Figure 4 illustrates the range of *K*-means clustering results using different values of *k*_cluster_ and *p*_axes_, with cluster fit evaluated using BIC scores (lower is more likely). For the *M* = 0.000 scenario, BIC scores rapidly decline and plateau at *k*_cluster_ = 5 in a relatively consistent way for all values of *p*_axes_. For the *M* = 0.004 scenario, the BIC scores exhibit an inflection at *k*_cluster_ = 5, although the shape of the BIC curves differ depending on *p*_axes_: whereas there is a plateauing pattern for *p*_axes_ = 10 and 20, there is an uptick pattern for *p*_axes_ = 40 and 80. For *M* = 0.400, there are also two different BIC curves: for *p*_axes_ = 10 and 20, BIC exhibits a steady declining pattern (inconsistent with *k* = 1), whereas for *p*_axes_ = 40 and 80, BIC exhibits a steady increasing pattern (consistent with *k* = 1). Notably, whilst *K*-means clustering should generally reflect the underlying *k*, different parameterisations can lead to different patterns in the test statistic curves. Values of *k*_infer_ obtained using *K*-means may vary across different parameterisations, *K*-means derived estimates of *k*_infer_ should be compared to expectations derived from PC screeplots and scatterplots to assess consistency across approaches.

**Figure 4.**
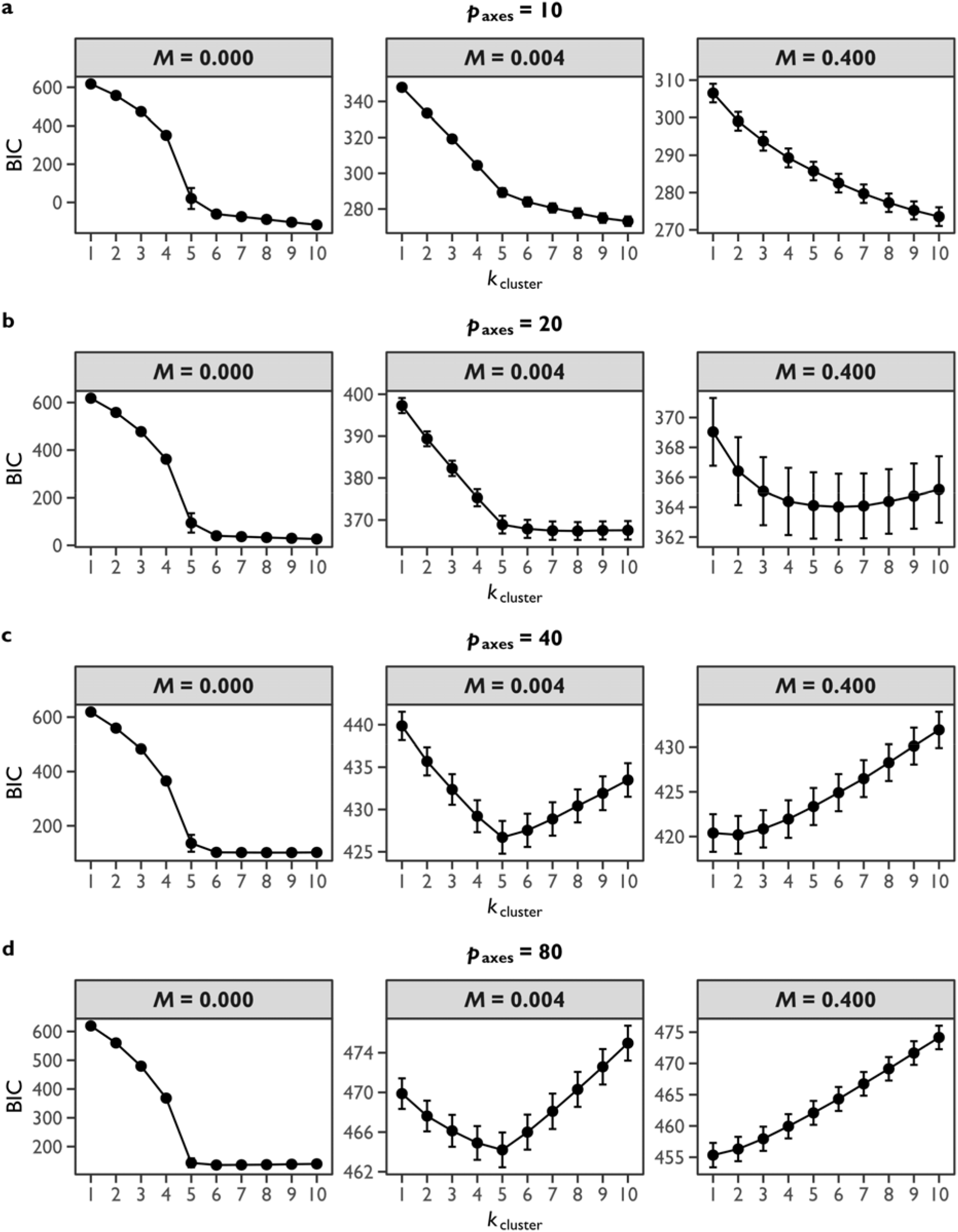
The inferred number of populations by *K*-means clustering, using the *replicate simulation dataset.* The *x*-axis is the number of clusters to divide samples among in *K*-means clustering, *k*_cluster_. The *y*-axis is the associated BIC score (lower values are more likely). Points represent the mean BIC across replicate simulations with 95% confidence intervals. Panels represent each of the simulated migration scenarios. Number of leading PC axes, *p*_axes_: (a) 10, (b) 20, (c) 40, and (d) 80.

Aside from the statistical principle that only the leading *k* – 1 PC axes capture population structure, these leading PC axes are also the only dimensions that are replicable. Replicability of PC axes in this context refers to recoverability of the same (or similar) eigenvectors across independent samples. Eigenvectors describe the relative contribution of each locus to each PC axis (the loadings). Loci contributing most to population structure should be consistent in the magnitude of their loadings. Where population structure exists, eigenvectors of the leading PC axes ≤ *k* – 1 are correlated across independent sample sets, whereas eigenvectors of PC axes ≥ *k* are not correlated across independent sample sets, as depicted in Figure 2b. However, the recoverability of eigenvectors depends on the magnitude of genetic differentiation. As genetic differentiation increases: (1) the intra-locus allelic correlation between individuals from the same population increases (Wright, 1949); (2) sampling bias is less likely to affect estimates of the inter-locus allelic covariances across all individuals; and (3) eigenvectors for the leading *k* – 1 PC axes will be more similar across independent sample sets. Compare, for example, the correlations between sample pair eigenvectors for the first four PC axes where *M* = 0.000 versus *M* = 0.004 versus *M* = 0.400, which show a decline with decreasing genetic differentiation. Moreover, in the *M* = 0.400 scenario, none of the eigenvectors have high correlations across independent sample sets because there is no population structure and no linear combinations of alleles that are repeatable. We should be concerned with this repeatability if we wish to use these PC axes as predictive variables, as discussed in the next section.

In summary, the analyses presented in this section illustrate how *k* can be inferred from a PCA of genotype data. Obtaining an inference of *k, k*_infer_, is important because only the leading *k* – 1 PC axes are biologically informative and exhibit repeatable linear combinations of loci across independent sample sets. PC axes ≤ *k* – 1 capture variation that is among populations, whereas PC axes ≥ *k* are associated with variation that is within populations. Obtaining *k*_infer_ is also useful because it can inform whether an initial *a priori* expectation of the number of populations, *k*_prior_, is valid.

## Suitable principal components for discriminant analysis

In this section, I will address the appropriate parameterisation of a DAPC. I will first discuss common practises in DAPC paramterisation and their (mis)alignment with the *k* – 1 limit of biologically informative PC axes. I will then provide a demonstration of how different parameterisations of a DAPC influence the solutions derived from its DA step. These analyses will highlight that only those PC axes ≤ *k* – 1 are suitable for DAPC. Inappropriate parameterisation of DAPC reduces the biological relevancy of the DA model and can give false impressions of population structure.

That we only expect the leading *k* – 1 PC axes to be biologically informative implies that the maximum value of *p*_axes_ should be *k* – 1, a *k* –1 criterion. However, typical DAPC parameterisation often involves choosing a value of *p*_axes_ that captures a certain proportion of the total genotypic variance, a *proportional variance criterion*. Another approach is to use the built-in functions from the *adegenet* package, xvalDapc and optim.a.score (Jombart, 2008; Jombart & Ahmed, 2011), to reduce and optimise the number of PC axes predictors, *p*’_axes_. The XVALDAPC function performs cross-validation for different numbers of *p*_axes_, inferring the optimal *p*’_axes_ as the one that produces the lowest mean squared error term. The optim.a.score function performs a reassignment analysis using true and randomised cluster identities to calculate an a-score for different numbers of *p*_axes_, inferring the optimal *p*’_axes_ as the one that produces the largest a-score.

Selecting *p*_axes_ to satisfy the *proportional variance criterion* is problematic. Unless there is very strong differentiation among populations, the value of *p*_axes_ required to capture a specific amount of genotypic variance will likely be much greater than *k* – 1. Additionally, the built-in optimisation functions have variable performance that may depend on the magnitude of genetic differentiation and the initial value of *P*_axes_ as illustrated in Figure 5. In my simulations, the xvalDapc function consistently returned a *p*’_axes_ value that was greater than the *k* – 1 = 4 expectation for *M* = 0.000 and *M* = 0.004 scenarios, and greater than the *k* – 1 = 0 expectation for *M* = 0.400, for all initiating values of *p*_axes_. The optim.a.score function tended to be more conservative but increasing migration rates and larger values of *p*_axes_ led to inflated estimates of *p*’_axes_.

**Figure 5.**
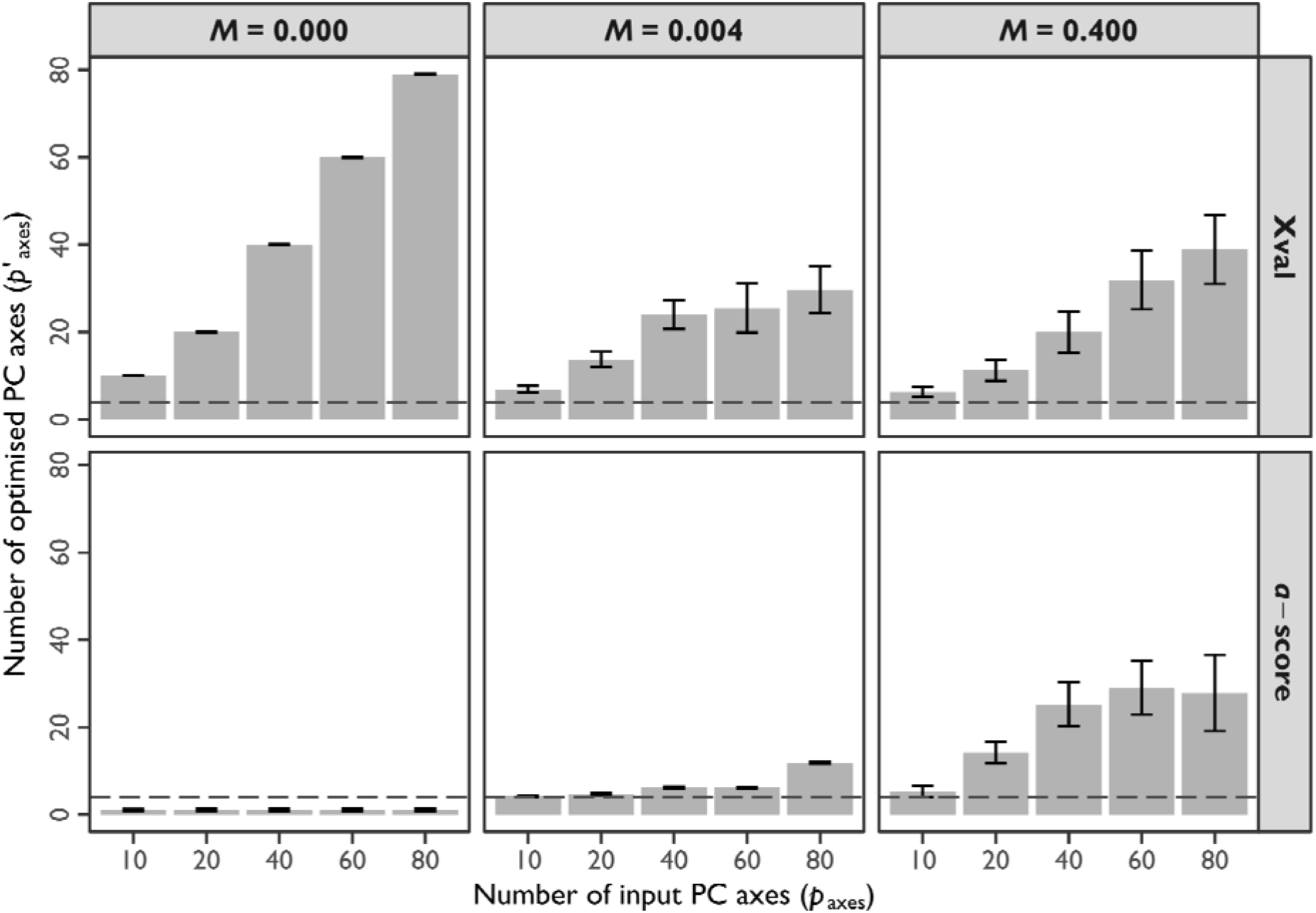
Optimisation of PC axes to fit DA models using the *replicate simulation dataset*. The *x*-axis is the number of PC axes used to initially fit the DA, *p*_axes_. The *y*-axis is the number of optimised PC axes, *p*’_axes_, using one of two optimisation algorithms available in the *adegenet* package. Panels represent combinations of optimisation algorithm (in rows) and migration scenario (in columns). Bars represent the mean *p*’_axes_ estimated across replicate simulations with 95% confidence intervals. The dashed lines demarcate *p*’_axes_ = 4.

Because population structure is only captured on the leading *k* – 1 PC axes, the inclusion of PC axes ≥ *k* does not provide additional information to discriminate among populations. We can think of the as the relative proportion of the variation in our set of predictor variables that is among versus within populations. Figure 6 depicts the mean variation among populations when using different values of *p*_axes_ to parameterise a DAPC with *k*_DA_ = 5 groups. For *M* = 0.000, irrespective of the value of *p*_axes_, the amount of variation among populations is always >95% because the first four PC axes explain virtually all the genotypic variance. However, in the *M* = 0.004 scenario, increasing *p*_axes_ reduces the variation among populations captured on these predictor variables. Finally, for *M* = 0.400, because there is no population structure, none of the PC axes capture among-population variation, and there is essentially a flat line across all values of *p*_axes_. When population structure exists, limiting *p*_axes_ to the leading *k* – 1 PC axes for DAPC parameterisation is more parsimonious, uses only biologically relevant predictors, and maximises the total variation among populations in the input predictor variables.

**Figure 6.**
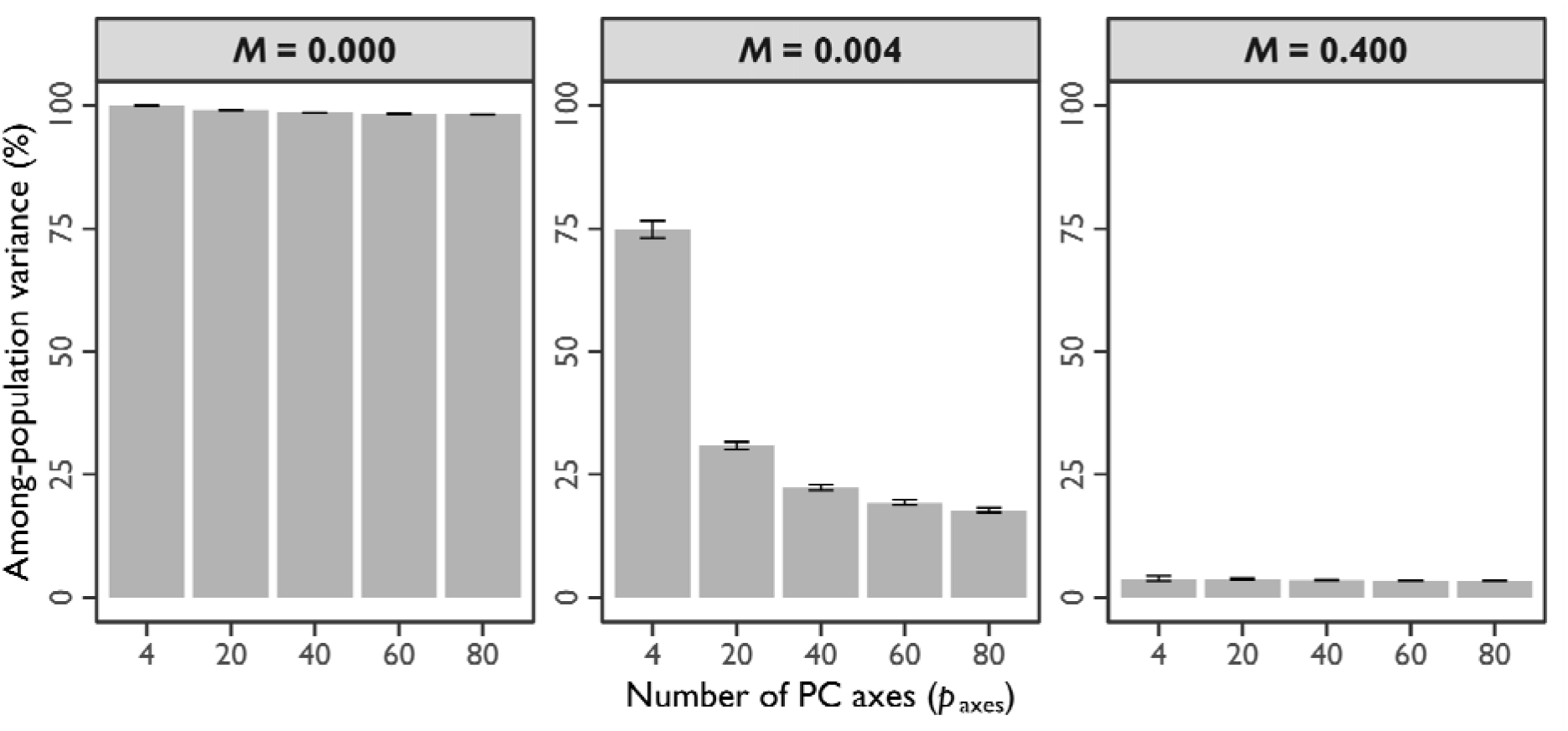
Average amount of among-population variance captured on PC axes using the *replicate simulation dataset*. The *x*-axis represents the total number of leading PC axes used in a DA, with *k*_DA_ = 5 groups. The *y*-axis represents the amount of variance (percent of total) that is among populations. Grey bars indicate the average across replicate simulations with 95% confidence intervals. Panels contain results for each migration scenario.

Although biologically informative PC axes should contribute most to the DA solution, as *p*_axes_ increases beyond *k* – 1, the DA model can be increasingly influenced by biologically uninformative predictors. Consider the contributions of PC axes in highly over-fit DA models using *p*_axes_ = 80 PC axes with *k*_DA_ = 5 groups. Figure 7 illustrates the average absolute loadings of the first 12 PC axes onto the discriminant functions describing four-dimensional LD space. For the *M* = 0.000 and *M* = 0.004, PC axes 1 to 4 (≤ *k* – 1) have consistently larger loadings relative to PC axes 5 and beyond (≥ *k*). Notably, for *M* = 0.000, each sequential LD axis largely corresponds to each PC axis in descending order from 1 to 4, as evidenced by the large absolute loadings. For *M* = 0.000, there is minimal within-population variation, so PC axes ≥ 5 have negligible contributions to the LD axes, and DA effectively returns the PCA results. For the *M* = 0.004 scenario, the first four PC axes combine in different ways to generate the LD axes. Unlike *M* = 0.000, DA does not simply return PCA results for *M* = 0.004, and the uninformative PC axes ≥ 5 have a larger influence on the final solutions. In the *M* = 0.400 scenario, *k* – 1 = 0, and the LD loadings are equivalent for all PC axes. For *M* = 0.400, all the variation is within populations, but DA derives a model of among-population differences despite none of the predictor PC axes capturing population structure. These data demonstrate that when genetic differentiation is large, the DA model is less likely to be influenced by the inclusion of many biologically uninformative PC axes. In contrast, when genetic differentiation is weak, inclusion of many biologically uninformative PC axes > *k* – 1 is more likely to influence the DA model. Furthermore, because DA is mechanistically a *hypothesis driven* method, a solution will be derived whether it is biologically meaningful or not.

**Figure 7.**
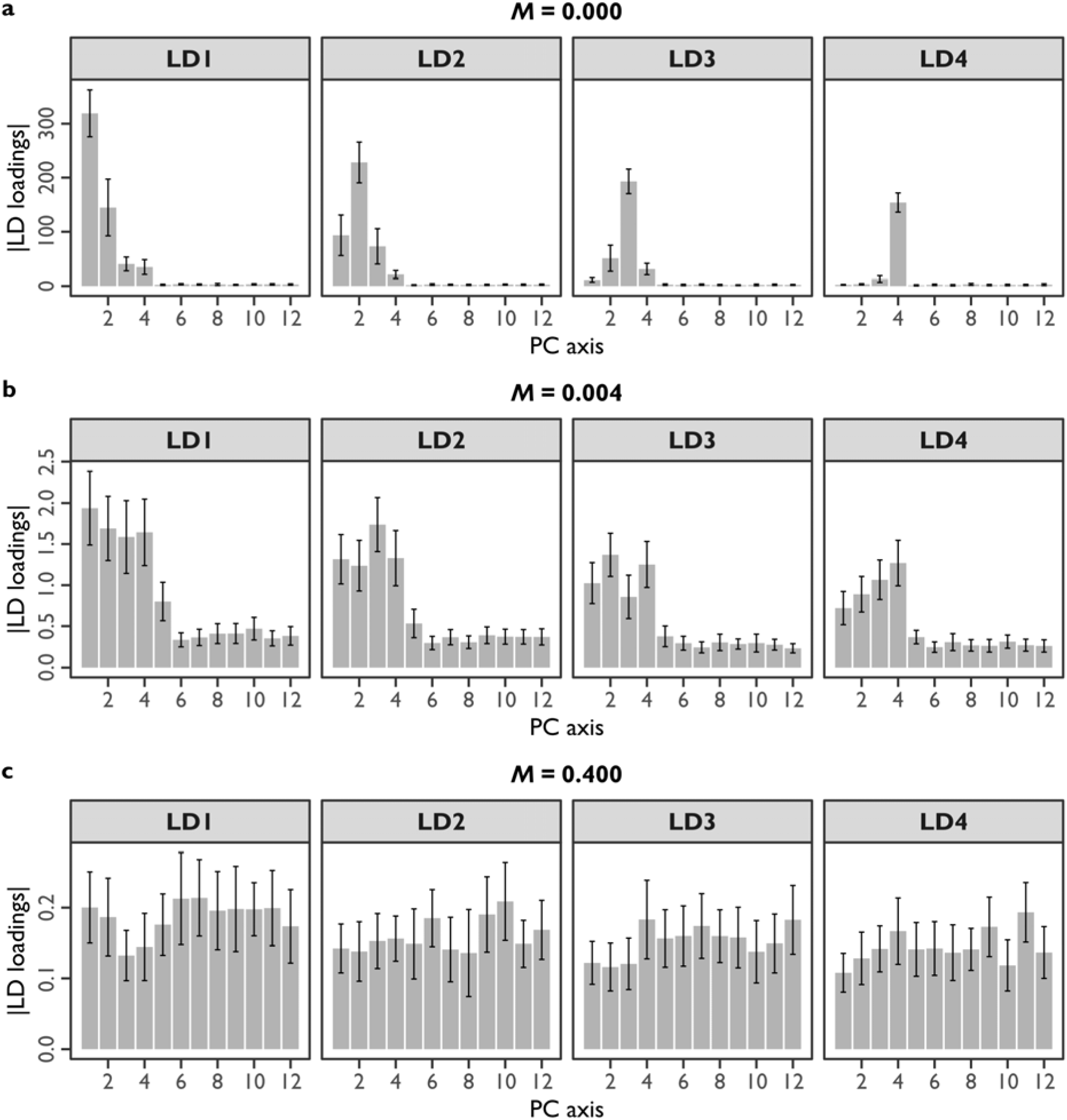
Average loadings of individual PC axes in DAs using the *replicate simulation dataset*. The *x*-axis represents the first 12 PC axes in DAs parameterised with *k*_DA_ = 5 and *p*_axes_ = 80 PC axes. The *y*-axis represents the absolute loadings (contributions to variation) of PC axes onto discriminant functions describing each of the four LD axes (panels). Grey bars indicate the average loadings across replicate simulations with 95% confidence intervals. Migration rates: (a) *M* = 0.000; (b) *M* = 0.004; and (c) *M* = 0.400.

By inspecting scatterplots of individuals projected into LD space, we can see how genetic differentiation and the choice of *k*_DA_ and *p*_axes_ interact in a DAPC (Figure 8). For all migration scenarios, increasing *p*_axes_ increases the separation of populations in LD space when *k*_DA_ = 5 groups; note the increasing scale of the LD axes and tighter clustering as *p*_axes_ increases. However, with respect to the general patterns inferred from the LD space projection, the effect of increasing *p*_axes_ » *k* – 1 is more dramatic when genetic differentiation is weak. For *M* = 0.000, regardless of whether *p*_axes_ = 4, 40 or 80 PC axes, we essentially recover the same projection of populations in LD space (compare Figures 8a, 8b, and 8c). For *M* = 0.004, the same general pattern emerges regardless of the value of *p*_axes_, but the perceived magnitude of this separation depends on *p*_axes_ (compare Figures 8d, 8e, and 8f). For *M* = 0.400, despite *k* = 1 and a lack of population structure, increasing *p*_axes_ gives the impression that populations can be reliably discriminated into discrete clusters. When using *p*_axes_ = 4 (Figure 8g), we retain a more realistic projection with negligible separation among populations in LD space. However, with *p*_axes_ = 40 and 80 (Figures 8h and 8i, respectively), there is increasing separation of populations in LD space. This perceived separation is not being driven by the inclusion of biologically relevant predictors of among-population differences, but instead, an over-fitting of the DA model to idiosyncrasies of the sample.

**Figure 8.**
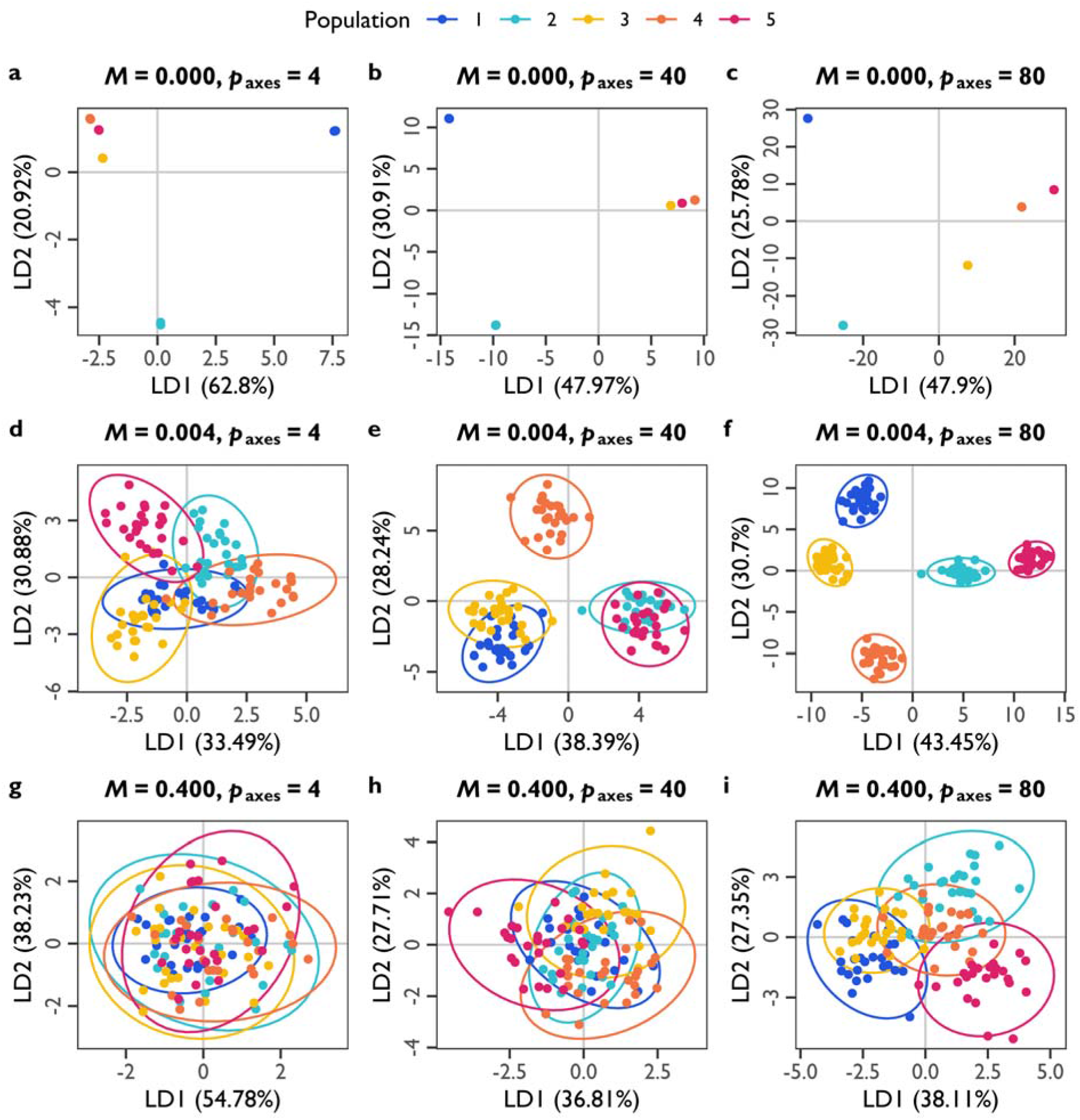
Exemplar scatterplots of individuals projected into LD space. The *x*-axis and *y*-axis represent the first and second LD axes, respectively, obtained from a DAPC parameterised with different numbers of PC axes, *p*_axes_, with *k*_DA_ = 5 groups. Numbers in parentheses indicate the amount of explained among-population variance for each LD axis (percent of total). Each point represents an individual, coloured by population (see legend). Migration rates: (a, b, c) *M* = 0.000; (d, e, f) *M* = 0.004; and (g, h, i) *M* = 0.400. Number of PC axes: (a, d, g) *p*_axes_ = 4; (b, e, h) *p*_axes_ = 40; and (c, f, i) *p*_axes_ = 80.

Because the DA model can be used for group assignment problems (see also the next section), I will use group assignment to further explore how different combinations of PC axes contribute (or do not contribute) to discrimination among populations. Our expectation is that when population structure exists, DA models including the first *k* – 1 PC axes will have better ability to assign individuals to their correct population, relative to DA models without these biologically informative PC axes. Figure 9 illustrates assignment analyses where a single *training* sample set was used to build a DA model with *k*_DA_ = 5 groups, and this model was then used to predict the populations of individuals in 19 *testing* sample sets. Different combinations of PC axes were used that either included the first four PC axes (1, 1–2, 1–3, 1–4, 1–40, and 1–80) or excluded them (5–40 and 5–80). Clearly, in the *M* = 0.000 and *M* = 0.004 scenarios (Figures 9a and 9b, respectively), the power to assign individuals to their correct population is being completely driven by the first four PC axes. Exclusion of the first four PC axes leads to poor correct assignment rates in these scenarios where population structure exists. For the *M* = 0.400 scenario (Figure 9c), there is no power to assign individuals, irrespective of the input p_axes_, because there are no biologically informative PC axes, *k* – 1 = 0.

**Figure 9.**
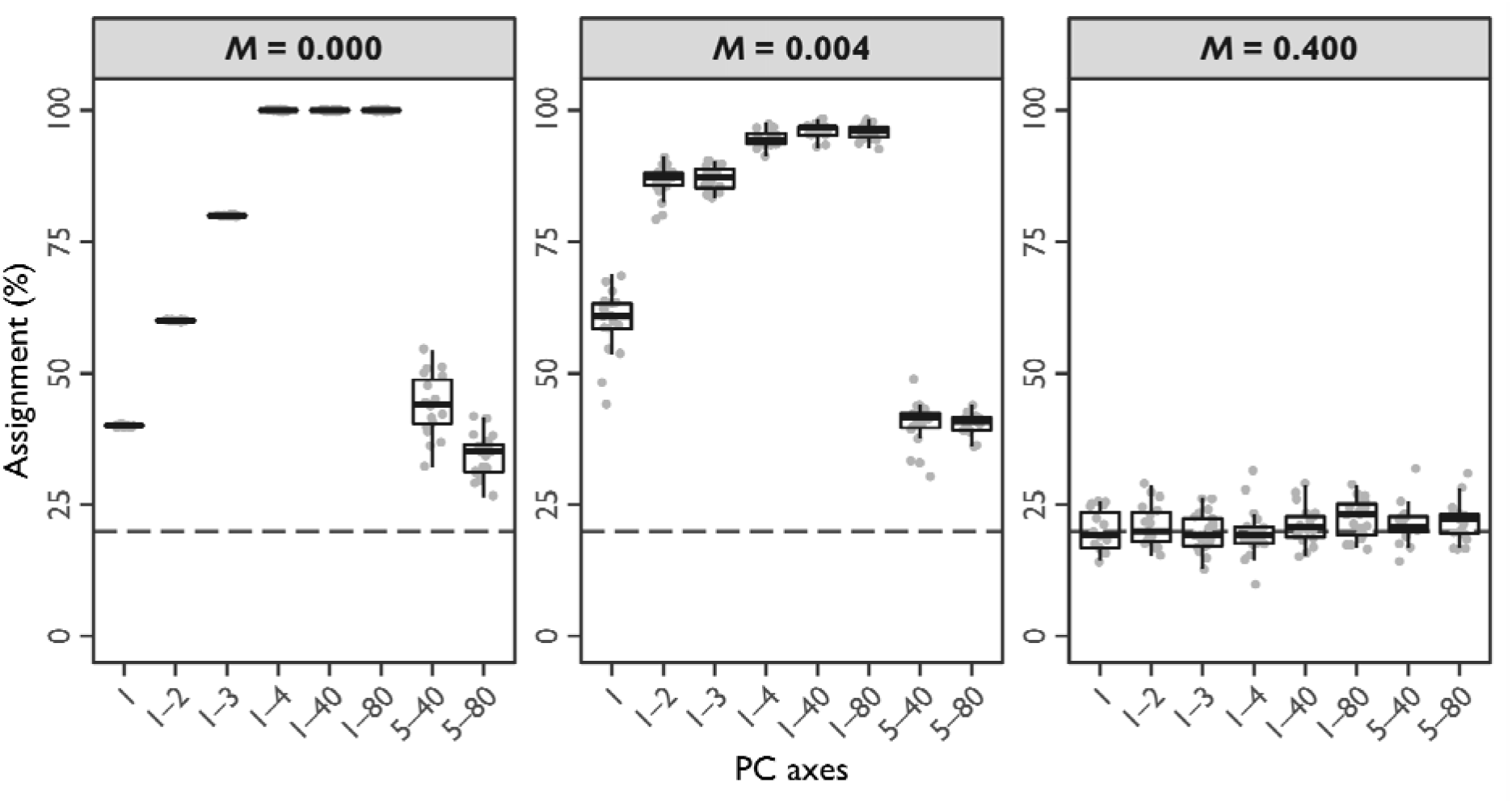
Correct assignment rates using DA models parameterised with different combinations of PC axes using the *singular simulation dataset*. DA models were constructed with a *training* sample set and used to assign populations in 19 *testing* sample sets, with a *k*_DA_ = 5. The *x*-axis represents the combinations of PC axes that either include the first four biologically informative *k* – 1 PC axes (1, 1–2, 1–3, 1–4, 1–40, and 180) or exclude them (5–40 and 5–80). The *y*-axis represents the percentage of correct assignment. Each grey point represents one of the *testing* sample sets. Boxplots summarise the distribution of correct assignment rates across *testing* sample sets. The dashed line demarcates a correct assignment rate of 20% (random assignment to any of the *k*_DA_ = 5 groups). Migration rates: (a) *M* = 0.000; (b) *M* = 0.004; and (c) *M* = 0.400.

The results from the assignment analyses (Figure 9) can be further interpreted by considering the recoverability of eigenvectors that were discussed above (Figure 2b). The linear combinations of loci that are described by eigenvectors are more repeatable across independent sample sets when there is greater genetic differentiation, but only for those *k* – 1 PC axes describing population structure (Figure 2b). In the context of a group assignment problem, whereas the first *k* – 1 PC axes provide repeatably useful predictors to assign populations, PC axes ≥ *k* are stochastic across different sample sets and are uninformative predictors for population assignment. The *M* = 0.400 scenario provides an extreme case where *k* = 1, there are no biologically informative PC axes, none of the eigenvectors are repeatable, and assignment of individuals to the five populations is not possible.

In summary, the analyses presented in this section demonstrate that only the leading *k* – 1 PC axes are suitable for DAPC of genotype data. The *hypothesis driven* nature of a DA requires appropriate parameterisation to ensure biologically meaningful results. Addition of PC axes greater than *k* – 1 does not add informative predictors to discriminate among *k*_DA_ groups in a DA. Parameterisation of *p*_axes_ ≫ *k* – 1 will result in a model being over-fit to the idiosyncrasies of the sample. However, the impact of this over-fitting the interpretation of population structure is likely to be greatest when genetic differentiation is weak. When genetic differentiation is weak, an over-fit DA model artificially inflates the perceived separation among groups in LD space, giving the impression that populations are more genetically discrete than they are. DAPC parameterised using only the leading *k* – 1 PC axes is maximised for the among-population variation in the predictor variables, is not influenced by the idiosyncrasies of uninformative predictors, and is generalisable to independent sample sets.

## Assessing model fit, interpreting population structure, and testing hypotheses

It is often unappreciated that when applying DAPC to genotype data a researcher is building a model that describes population structure. The results of DAPC are almost exclusively interpreted through visualising the projection of samples into LD space, but rarely are assessments of DA model fit reported. As a *hypothesis driven* method, DA is effectively testing the hypothesis that there are significant differences among defined groups. However, whether a researcher can interpret their DAPC results as a true test of this hypothesis depends on whether they defined groups *a priori* or *de novo.* In this section, I outline how to assess the fit of a DA model and how assessing model fit can aid interpretation of population structure. I also discuss considerations for interpreting DAPC results relative to a researcher’s goals and their rationale behind group designations.

Two approaches that can be used to assess the fit of a DA model are leave-one-out cross-validation (for small sample sizes) and training-testing partitioning (for large sample sizes). Leave-one-out cross-validation involves performing a series of *n* iterative model fits for all *i* = 1 to *i* = *n* individuals in a dataset. For each *i*^th^ iteration, the *i*^th^ individual is withheld from model fitting, the model is fitted with the remaining *n* – 1 individuals, and the group identity of the withheld i_th_ individual is predicted using the fitted model. Training-testing partitioning requires random division of a dataset to create a training partition of 70–80% of individuals to fit the model, which can then be used to predict the groups of the 20–30% withheld individuals in the testing partition. Correct assignment rates can then be calculated from model predictions. A global correct assignment rate can be used to assess the model’s fit overall, providing an indication of how well any individual can be correctly assigned to their population and the discreteness of populations most generally. More nuanced perspectives can be attained by calculating correct assignment rates for each population and by examining patterns of incorrect assignment (see for example, Table S1). Correct assignment of individuals from certain populations might be more (or less) likely, indicating that the model is better (or worse) at discriminating these populations. Examining predictions of incorrectly assigned individuals can be used to identify populations that share genetic variation and have less discrete genetic boundaries. Correct assignment rates of ≥90% indicate that the DA model can discriminate individuals with high confidence, although an exact threshold for a “good” model is arbitrary.

It is important to note that whilst a researcher can always use their DAPC to interpret population structure, they should not necessarily use their DAPC to test the hypothesis that there are significant genetic differences among populations. This may seem counter intuitive given my description of DA as a *hypothesis driven* method. However, the distinction depends on a researcher’s goals and the rationale behind their choice of *k*_DA_. Mechanistically, *k*_DA_ represents the tested hypothesis within the DA framework. But if the choice of *k*_DA_ is influenced by the data, then this is not a true hypothesis test. For example, say a researcher samples individuals of an organism at five different locations, but upon examination of the genotype data, discovers that some of the locations are genetically indistinguishable, such that there are only three genetically distinct groups. The researcher might then be interested in discriminating among these inferred genetic groups, performing a DAPC with *k*_DA_ = *k*_infer_ = 3 groups and p_axes_ = *k*_infer_ – 1 = 2 PC axes as predictors. Whilst the researcher can use their DA model to interpret population structure (the relationships and patterns among inferred groups), they should not use this model to conclude that there are significant genetic differences among these inferred groups. To do so would be circular because groups were identified *de novo,* that is, *after* the researcher observed the genotype data (Meirmans, 2015).

If instead a researcher defines groups *before* they observe their genotype data, then results from a DAPC can be truly considered as a hypothesis test of genetic differences among groups. For example, say a researcher samples individuals of an organism at five different locations, and they believe *a priori* that individuals from those locations comprise discrete populations. The researcher could then perform a DAPC with *k*_DA_ = *k*_prior_= 5 groups to test whether it is possible to discriminate among these locations as evidence of population structure. To determine *p*_axes_, the researcher would then need to estimate the number of effective populations, *k*_infer_, to identify the number of leading PC axes that are biological informative, using *p*_axes_ = *k*_infer_– 1 PC axes as predictors. The value of *k*_infer_ may be the same as the researcher’s expectations (*k*_prior_ = *k*_infer_), or it may differ from their expectations (*k*_prior_ > *k*_infer_, or *k*_prior_ = *k*_infer_). Regardless, because estimation of *k*_infer_ does not inform the choice of *k*_DA_, there is no issue with circularity and the researcher is performing a true hypothesis test when fitting their DA model.

In summary, it is crucial that researchers are clear in advance about their goals when applying DAPC to genotype data. Do they want to test how genetic variation is organised among populations defined *a priori* (*k*_DA_ = *k*_prior_), or do they want to identify populations *de novo* and examine the genetic variation among these inferred groups (*k*_DA_= *k*_infer_)? These different goals dictate how a researcher should parameterise and interpret their DAPC, and clearly communicating these goals will allow readers to better understand a researcher’s rationale and results. Researchers should more routinely provide an assessment of their DA model fit, which can be used to evaluate how well their model discriminates among populations and aid interpretation of population structure. Testing the hypothesis that significant genetic differences exist among population with a DA model should only be considered when a researcher has well-defined *a priori* expectations.

## Concluding remarks

As a *hypothesis free* method, PCA is useful for visualising the inherent structure present in a dataset. Contrastingly, DA is a *hypothesis driven* method and generates a visualisation that maximises the differences among defined groups. The combination of these methods into a DAPC of genotype data has become very popular. DAPC provides a fast and simple method to summarise population structure and to perform population assignment. The ease at which DAPC can be applied to genotype data in R, and its lack of demographic assumptions, make it a wonderfully flexible analysis. Yet paucity of clear best practise guidelines has made the implementation, interpretation, and reporting of DAPC results inconsistent (Miller et al., 2020). DAPC should not be considered a method for “finding population structure”, but instead, a method used to model the genetic differences among groups that a researcher is interested in. As I have demonstrated, parameterisation of DAPC matters, and the guidelines I present here will help promote standardisation of DAPC in future studies.

My work demonstrates that only the leading *k* – 1 biologically informative PC axes should be used as predictors in a DAPC of genotype data. I show that this *k* – 1 criterion sets a deterministic value for *p*_axes_ specification and is more suitable than the commonly used *proportional variance criterion.* Inclusion of many biologically uninformative PC axes as predictors of genetic differences among *k*_DA_ groups in DA can lead to over-discrimination of these groups in LD space. Projection of individuals into LD space is typically a focal point when interpreting DAPC results. Hence, artificially inflated separation among groups is problematic because the perceived population structure can appear greater than that present in the genotype dataset.

Whereas other population genetic methods implementing PCA perform a selection step to limit analyses to biologically informative PC axes (for example: Conomos, Reiner, Weir, & Thornton, 2016; Luu, Bazin, & Blum, 2017; Meisner & Albrechtsen, 2018), such considerations have never been discussed in the context of performing a DAPC, to the best of my knowledge. This is surprising given existing works describing the link between PCA and the genetic relationships among individuals and populations (McVean, 2009; Patterson et al., 2006; Peter, 2022). There are many ways to estimate the number of biologically informative PC axes (Cattell, 1966; Jackson, 1993; Peres-Neto et al., 2005). For genotype data, Patterson et al. (2006) provided a test based on expectations under a Tracy-Widom distribution to statistically infer *k*. Notwithstanding, careful examination of PC screeplots and scatterplots provide a simple visual way to infer *k*, and this inference can be corroborated with estimates from K-means clustering, a standard approach used alongside DAPC for inferring genetic groups *de novo*.

Assessing the fit of the DA model should be routine practise when performing a DAPC of genotype data. The rarity of such practise is perhaps due to general misunderstanding among researchers that they are fitting a model of genetic differences among groups with this method. When interpreting the outputs of their DA model, a researcher should be conscious of their study goals and their rationale behind defining *k*_DA_ groups. The DA model can only be used to test the hypothesis of significant genetic differences among groups when *k*_DA_ is defined a priori, *k*_DA_, = *k*_prior_. When DA is used to model differences among *de novo* inferred groups, *k*_DA_ = *k*_infer_, interpretation should be limited to patterns of population structure to avoid circularity.

Irrespective of whether a researcher’s DAPC entails a test of an *a priori* expectation or the study of *de novo* inferred groups, an estimate of the likely *k* is required for choosing an appropriate *p*_axes_. Under more complex demographic scenarios, it may be challenging to estimate *k*_infer_ when populations do not form easily discernible groups (see for illustration, Figure S4). Nonetheless, the expectations of a *k* – 1 limit on the number of biologically informative PC axes will still hold, and a researcher should do their best to make an appropriate judgement. A small discrepancy between *k* and *k*_infer_ is unlikely to be cause notable problems in the DA model fit, the point being that *p*_axes_ ≫ *k* – 1 is more problematic and should be avoided.

Suggestions for future population genetic studies include:

1. Clearly state whether the purpose of implementing a DAPC of genotype data is to test genetic differences among *a priori* defined populations (*k*_DA_ = *k*_prior_) or to study variation among *de novo* inferred populations (*k*_DA_ = *k*_infer_).
2. If k-means clustering is used to obtain *k*_infer_, check that results are consistent across different parameterisations, and that these results are also consistent with expectations derived from visual inspection of PC screeplots and scatterplots.
3. Use the *k* – 1 criterion to select an appropriate *p*_axes_ for DAPC. Because *k* is never known, this involves setting *p*_axes_ = *k*_infer_ –1 to retain only the leading biologically informative axes for the putative *k*_infer_ effective populations.
4. Assess the DA model fit using leave-one-out cross-validation (for small sample sizes) or training-testing partitioning (for larger sample sizes). Report global correct assignment rates and relative proportions of correct and incorrect assignment among populations.
5. Check that *k*_infer_ is not artificially inflated, for example, by the presence of highly correlated genotypes in sex chromosomes or inversions, or by the presence of family structure within populations. Sex-linked loci should be removed for analyses comprising pooled sexes because they will polarise males and females. Loci within inversions can either be removed, or inversion genotypes can be recoded as a single “super locus”. Family structure represents non-random sampling of populations but may be unavoidable in small populations. Being aware of the effects of family structure is important for interpreting a larger number of groups relative to *a priori* expectations. Additionally, if groups are being defined *de novo,* then it may be nonsensical to discriminate among all inferred groups if a researcher is interested in differences among putative populations.
6. When *k*_infer_ = 2, there is just a single dimension that is biologically informative: the first PC axis. A DAPC can still be fit with a single predictor variable, although it defeats the purpose of using a multivariate analysis and is equivalent to a univariate analysis.

It is prudent to reflect on whether a DAPC is necessary for the specific objectives of a study. If a researcher simply wants to visualise population structure, a PCA alone is perfectly sufficient and does not require allocation of group membership. However, when there are many putative populations, visualising just a few PC axes might be inadequate to fully appreciate the complexity of population structure. In such a case, DAPC may be helpful for summarising variation across many relevant PC axes and to further reduce dimensionality. The real value of a DAPC comes when we take advantage of the predictive model produced by the DA. These predictive models can be used in assignment problems to identify source populations of recent migrants or to infer the parental populations in admixture events. The individual contributions of loci to variation among populations could also be exploited to identify candidates for divergent selection. Irrespective of the goal, limiting *p*_axes_ to the leading *k* – 1 PC axes will produce a DA model fit that is more parsimonious and captures maximal among-population variation from biologically relevant predictor variables. An appropriately parameterised DA model is less likely to suffer from unintended interpretations of population structure and is more generalisable to individuals from independent samples sets.

## Acknowledgements

I thank Cynthia Riginos and Tom Schmidt for providing feedback on this manuscript. This piece benefitted from the thoughtful critiques of two anonymous reviewers, to whom I am grateful.

## Data accessibility statement

The data and scripts associated with this manuscript have been deposited into *Dryad* (Thia, 2022).

## R packages

*adegent* (Jombart, 2008; Jombart & Ahmed, 2011)

*data.table* (Dowle & Srinivasan, 2019)

*doParallel* (Microsoft Corporatin & Steve Weston, 2019)

*extrafont* (Chang, 2014)

*genomalicious* (Thia & Riginos, 2019)

*ggpubr* (Kassambara, 2020)

*ggtree* (Yu, Smith, Zhu, Guan, & Lam, 2017)

*MASS* (Venables & Ripley, 2002)

*tidyverse* (Wickham et al., 2019)

## Supplementary Information: Analytical methods

### *F*_ST_ calculations

Estimates of Weir and Cockerham’s *F*_ST_ were performed using the fstWC_genos function from the R package, *genomalicious*.

### Eigenvector correlations

Eigenvector correlations were determined between pairs of independent sample sets in the *singular simulation dataset*. For each sample set, I performed a PCA on the allelic covariances using R’s procomp function, with the arguments “scale=FALSE, center=TRUE”. For each *i*^th^ eigenvector from 1–10, I calculated the Pearson correlation between each *j*^th^ sample set pair, using R’s cor function. This allowed me to determine how similar each *i*^th^ eigenvector was among the different sample sets.

### Discriminant analysis of principal components

Multiple DAPCs were performed on independent simulations from the *replicate simulation dataset*. The *genomalicious* function adegent_DT2genelight was used to convert genotype data tables into *adegenet* ‘genlight’ objects. The dapc function in *adegenet* was then used to fit DAPCs using *p*_axes_ = 4, 20, 40, 60, and 80 PC axes.

### Among-population variation

The proportion of among-population variation captured by each value of *p*_axes_ in the *replicate simulation datasets* was calculated by performing a multivariate analysis of variance (MANOVA). I used R’s MANOVA function to fit the model:

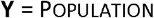

Where **Y** is a matrix of PC axes and Population is a categorical factor of population identity. I used the summary function on the ‘manova’ object to extract the sum of squares and cross products matrices for the hypothesis and error term, **H** and **E**, respectively. The proportion of among-population variation was then calculated as:

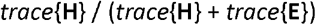

Where *trace*{} indicates the trace of the associated matrices.

### *K*–means and optimisation of input PC axes

For *p*_axes_ = 10, 20, 40, 60 and 80 PC axes, I estimated *k*_infer_ and tested the performance of optimisation algorithms to derive an optimised number of PC axes, *p*’_axes_. These analyses were conducted on simulations from the replicate simulation dataset, *K*-means clustering was performed using R’s kmeans. For each value of *p*_axes_, I tested support for *k*_cluster_ = 1 to 10 clusters. A BIC score was calculated as support for each value of *k*. Optimisation to derive *p*’_axes_ was conducted using the xvalDapc and optim.a.score functions from the *adegenet* package. For each function and input *p*_axes_ combination, I tested all possible 1 to *p*_axes_ number of PC axes, each with 50 iterations.

### Population assignment

Population assignment analyses were conducted using the *singular simulation dataset.* One of the independent sample sets was used as a *training* sample set, with the remaining 19 sample sets used as *testing* sample sets. PCA of allelic covariances in the *training* sample set was conducted using the PRCOMP function with the arguments “scale=FALSE, center=TRUE”. A series of DAPC models were manually fit using different combinations of PC axes from the *training* sample sets: 1, 1–2, 1–3, 1–4, 1–40, 1–80, 5–40, and 5–80 PC axes. The lda function from the *MASS* package was used to perform a DA using these different combinations of PC axes. To test the discriminatory power of this model, individuals from each *testing* sample set were first projected into the same PC space as the *training* sample set, using eigenvectors from the *training* sample set PCA. Next, the DA model constructed on the *training* sample set was used to take the PC axes of the *testing* sample sets and project them into LD space. Finally, population assignment was performed using R’s predict function.

### Supplemental simulations of more complex demographic scenarios

To illustrate the *k* – 1 limit of biologically informative PC axes under more complex demographic scenarios, I modelled isolation-by-distance and hierarchical structure using *fastsimcoal2.* Like the core simulations, I modelled five populations following a series of sequential divergence events (Figure S1), with each population containing 500 diploid individuals, and genome with a length of 20 Mb, a recombination rate of 1e–8, and a mutation rate of 2e–9. At the end of the simulation, I sampled 25 diploids from each population.

For the isolation-by-distance simulation, I modelled migration between adjacent populations in a sequential array: 1, 2, 3, 4, 5. Adjacent populations were connected by a pairwise migration rate of *m* = 0.005. Therefore, populations with two neighbouring populations (2, 3 and 4) had a total migration rate of *M* = 0.010, and those at the ends of the population array (1 and 5) had a total migration rate of *M* = 0.005.

For the hierarchical structure simulation, I modelled two levels of migration: within and between groups. Populations were split into two groups, with populations 1 and 2 forming group 1, and populations 3, 4 and 5 forming group 2. Pairwise migration rates were *m* = 0.0025 between populations within regions, whereas populations between regions were connected by a pairwise migration rate of *m* = 0.0005.

SNP genotypes were imported into R. A PCA of allelic covariances was performed for each demographic scenarios using all SNPs. *K*-means clustering was used to assess the number of effective populations from *k*_cluster_ = 1 to 10, with *p*_axes_ = 80. BIC scores were calculated as the test statistic for each value of *k*.

### Supplementary Information: Figures

**Figure S1.**
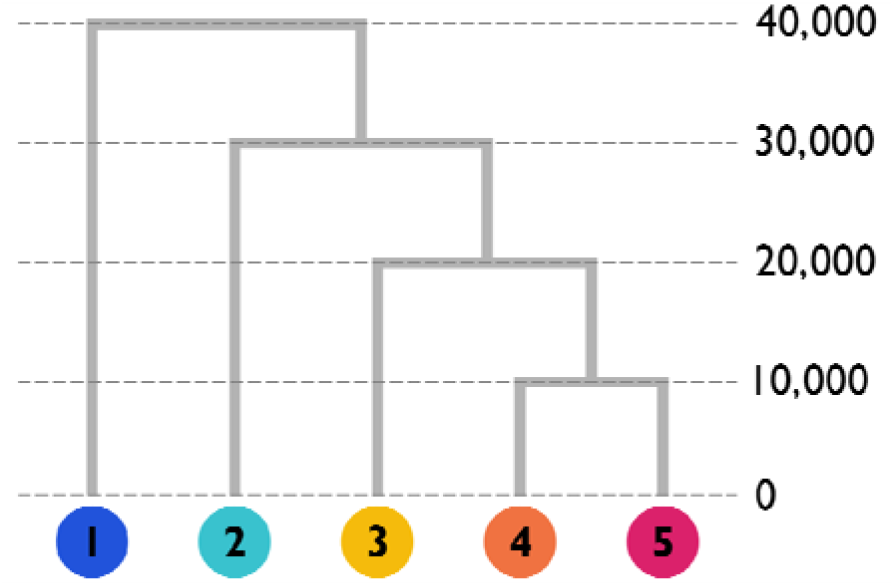
Simulated population divergence times. Each population (1–5) was comprised of 500 diploid individuals. Annotated nodes demarcate the number of generations in the past for each divergence event.

**Figure S2.**
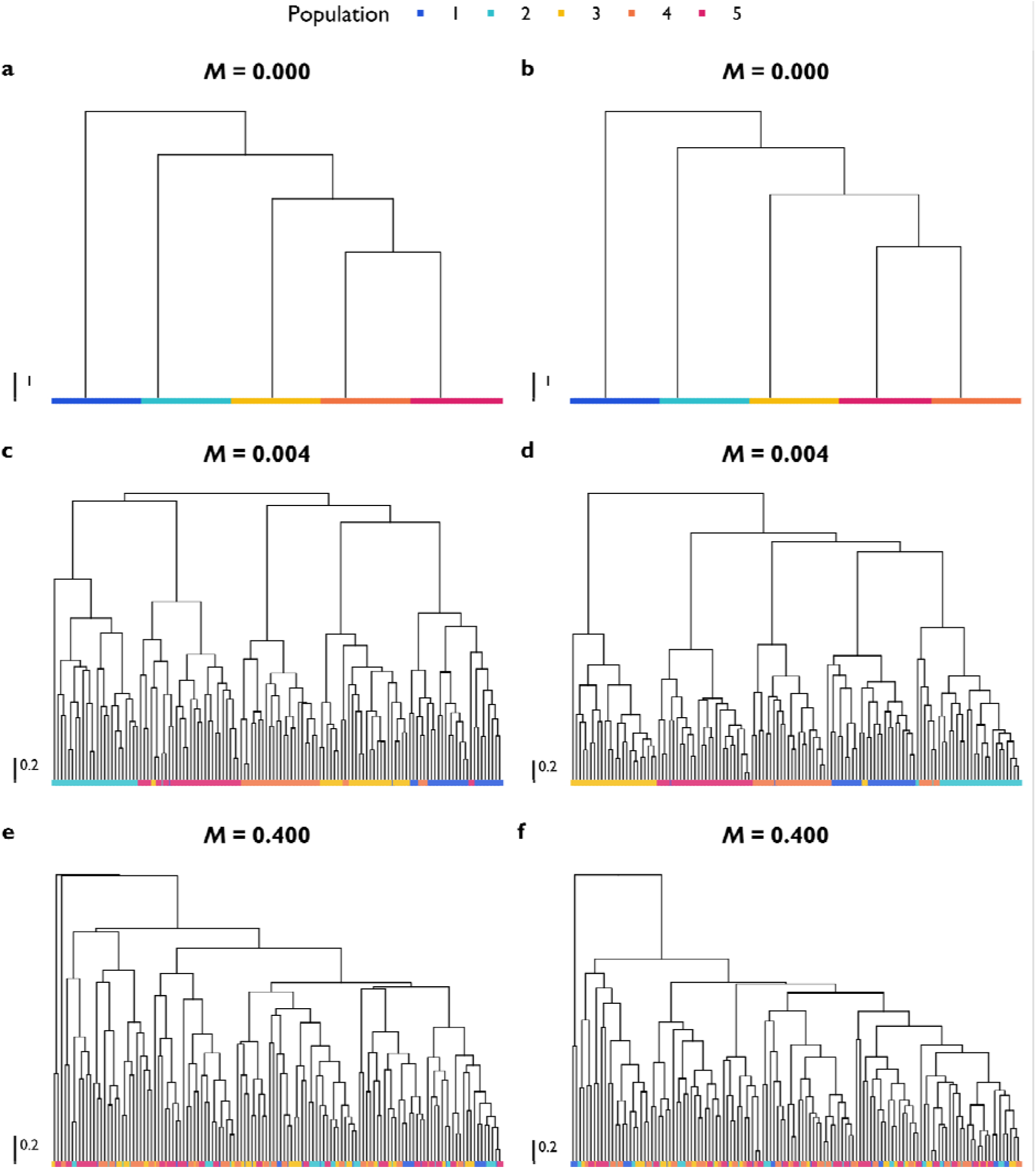
Hierarchical clustering of simulated samples in PC space from the *replicate simulation dataset.* Euclidean distance was estimated between all pairs of individuals using PC axes 1–4. Distance matrices were subjected to hierarchical clustering using the UPGMA method (unweighted pair group method with arithmetic mean) in R’s HCLUST function from the *stats* package. Branch lengths depict the Euclidean distance among individuals (see scalebar). Tips represent individuals, coloured by population (see legend). Examples of replicate simulations for each migration scenario: (a,b) *M* = 0.000; (c,d) *M* = 0.004; and (e,f) *M* = 0.400.

**Figure S3.**
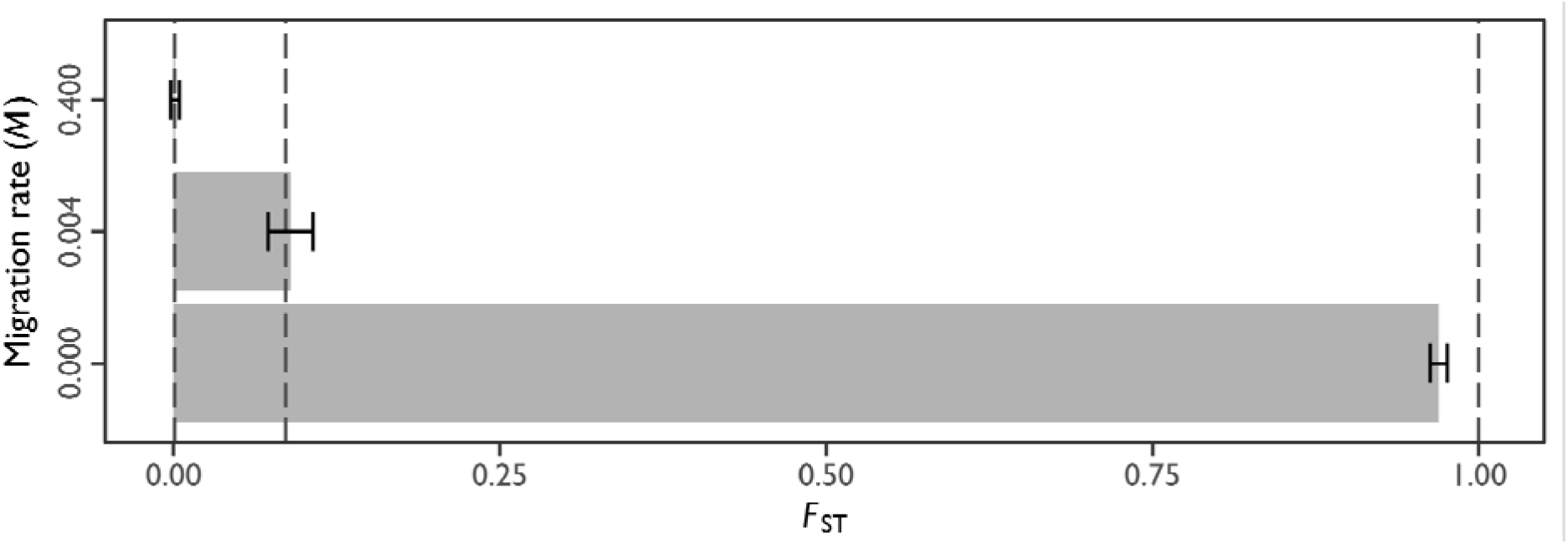
*F*_ST_ among populations in the *replicate simulation dataset.* The y-axis is the simulated migration rate and the *x*-axis is the estimated *F*_ST_. Grey bars depict the mean *F*_ST_ across replicate simulations with 95% confidence intervals. Dashed lines demarcate the expected *F*_ST_ values at each migration rate (from left to right): *F*_ST_ = 0.0009, 0.09, and 0.99, for *M* = 0.400, 0.004, and 0.000, respectively.

**Figure S4.**
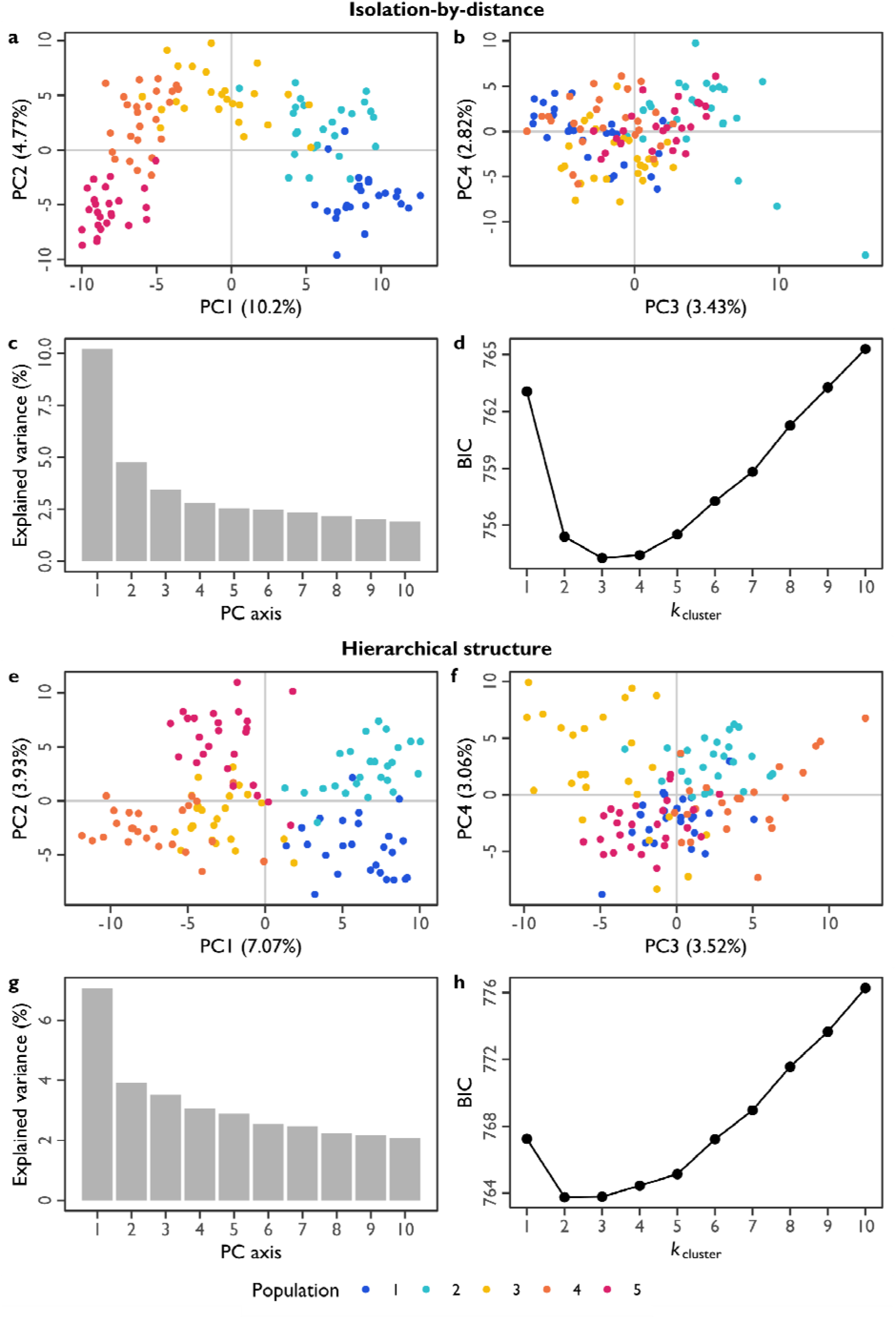
Population structure under more complex demographic scenarios, (a–d) Isolation-by-distance. (e–h) Hierarchical structure, (a, b, e, f) PC scatter plots. The *x*-axis and *y*-axis, respectively, represent different pairs of PC axes. Numbers in parentheses indicate the amount of explained genotypic variance captured by each PC axis (percent of total). Each point represents an individual, coloured by population (see legend), (c, g) Scree plots of explained variance. The *x*-axis represents the first 10 PC axes. The *y*-axis is the amount of explained genotypic variance (percent of total) ascribed to each PC axis, (d, h) Predicted number of populations using *K*-means. The *x*-axis is the inferred number of clusters using *p*_axes_ = 80 PC axes. The *y*-axis is the associated BIC score (lower values are more likely).

### Supplementary Information: Tables

**Table S1.** Assignment proportions using training-testing dataset partitions under different migration rates across samples from the *singular simulation dataset*. The true populations are in the rows, and the predicted populations are in the columns. The diagonal reflects the correct assignment rate.

| (a) $M = 0.000$ | | | | | |
| --- | --- | --- | --- | --- | --- |
|  | 1 | 2 | 3 | 4 | 5 |
| 1 | 1.000 | 0.000 | 0.000 | 0.000 | 0.000 |
| 2 | 0.000 | 1.000 | 0.000 | 0.000 | 0.000 |
| 3 | 0.000 | 0.000 | 1.000 | 0.000 | 0.000 |
| 4 | 0.000 | 0.000 | 0.000 | 1.000 | 0.000 |
| 5 | 0.000 | 0.000 | 0.000 | 0.000 | 1.000 |
| (b) $M = 0.004$ | | | | | |
|  | 1 | 2 | 3 | 4 | 5 |
| 1 | 0.947 | 0.004 | 0.002 | 0.002 | 0.044 |
| 2 | 0.006 | 0.966 | 0.006 | 0.004 | 0.017 |
| 3 | 0.023 | 0.002 | 0.943 | 0.000 | 0.032 |
| 4 | 0.053 | 0.029 | 0.008 | 0.893 | 0.017 |
| 5 | 0.004 | 0.008 | 0.002 | 0.000 | 0.985 |
| (c) $M = 0.400$ | | | | | |
|  | 1 | 2 | 3 | 4 | 5 |
| 1 | 0.320 | 0.242 | 0.069 | 0.331 | 0.038 |
| 2 | 0.284 | 0.206 | 0.088 | 0.381 | 0.040 |
| 3 | 0.324 | 0.219 | 0.088 | 0.318 | 0.051 |
| 4 | 0.364 | 0.208 | 0.065 | 0.324 | 0.038 |
| 5 | 0.347 | 0.198 | 0.082 | 0.333 | 0.040 |

